# Pavlovian cue-evoked alcohol seeking is disrupted by ventral pallidal inhibition

**DOI:** 10.1101/2024.03.14.585064

**Authors:** Jocelyn M. Richard, Anne Armstrong, Bailey Newell, Preethi Muruganandan, Patricia H. Janak, Benjamin T. Saunders

## Abstract

Cues paired with alcohol can be potent drivers of craving, alcohol-seeking, consumption, and relapse. While the ventral pallidum is implicated in appetitive and consummatory responses across several reward classes and types of behaviors, its role in behavioral responses to Pavlovian alcohol cues has not previously been established. Here, we tested the impact of optogenetic inhibition of ventral pallidum on Pavlovian-conditioned alcohol-seeking in male Long Evans rats. Rats underwent Pavlovian conditioning with an auditory cue predicting alcohol delivery to a reward port and a control cue predicting no alcohol delivery, until they consistently entered the reward port more during the alcohol cue than the control cue. We then tested the within-session effects of optogenetic inhibition during 50% of cue presentations. We found that optogenetic inhibition of ventral pallidum during the alcohol cue reduced port entry likelihood and time spent in the port, and increased port entry latency. Overall, these results suggest that normal ventral pallidum activity is necessary for Pavlovian alcohol-seeking.

## INTRODUCTION

Conditioned cues paired with alcohol and other drugs can contribute to misuse of these substances in a variety of ways (Stewart et al., 1984; Shaham et al., 2003; Robinson and Berridge, 2004; Everitt and Robbins, 2005; Valyear et al., 2017; Poisson et al., 2021). Alcohol cues can elicit or exacerbate relapse to alcohol seeking and consumption (Niaura et al., 1988; Garland et al., 2012; Garbusow et al., 2014) and lead to escalated alcohol intake (Stein et al., 2000; Koordeman et al., 2011; Jones et al., 2013). These effects have been attributed to the ability of conditioned cues to produce craving and physiological arousal, to impair inhibitory control, and to reduce cognitive resources (Sayette and Hufford, 1994; Tiffany and Carter, 1998; de Wit, 2009; Field and Jones, 2017). The neural mechanisms underlying these processes, particularly in response to alcohol cues, remain to be established.

The ventral pallidum (VP) is a basal forebrain structure implicated in appetitive motivation, consummatory behavior and positive affective responses, among other behaviors (Smith et al., 2009; Root et al., 2015; Soares-Cunha and Heinsbroek, 2023). It has been suggested to act as part of “final common pathway” for relapse to drug and alcohol seeking, including in response to cues (Kalivas and Volkow, 2005; Saunders et al., 2015; Kupchik and Prasad, 2021). Inhibition of VP disrupts behavior across several relapse models (McFarland and Kalivas, 2001; McFarland et al., 2004; Tang et al., 2005; Mahler et al., 2014; Farrell et al., 2019, 2022). In models of relapse to alcohol seeking, VP circuits have been implicated in context-induced renewal of alcohol seeking (Perry and McNally, 2013; Prasad and McNally, 2016; Prasad et al., 2020), but the causal contributions of VP activity to behavioral responses to Pavlovian alcohol cues, in reinstatement models or otherwise, have not been established. This may be due in part to the more limited effectiveness of discrete response-contingent cues to drive relapse-like behavior for alcohol reward (Tsiang and Janak, 2006; Richard and Fields, 2016).

Prior work from our group and others has indicated that VP activity both encodes and causally contributes to the likelihood of sucrose seeking in response to Pavlovian cues (Tindell et al., 2004, 2005; Richard et al., 2018; Stephenson-Jones et al., 2020) and the likelihood and vigor of sucrose seeking in response to discriminative stimuli (Richard et al., 2016, 2018; Lederman et al., 2021; Scott et al., 2023). Notably, we also found that the activity of VP neurons and ensembles encodes Pavlovian alcohol cues rather poorly, relative to Pavlovian sucrose cues (Ottenheimer et al., 2019).

To build on this past work, we investigated the impact of VP inhibition on Pavlovian-cue conditioned alcohol-seeking behavior, using a temporally specific optogenetic approach. Male Long Evans rats underwent Pavlovian conditioning with auditory cues predicting delivery of alcohol or no reward, followed by test sessions where VP neurons were inhibited on 50% of cue presentations. Despite the relatively poor encoding of Pavlovian alcohol cues by VP neurons (Ottenheimer et al., 2019), we find that VP activity during these cues is necessary for normal conditioned alcohol-seeking responses. Specifically, VP inhibition during the alcohol cue made rats less likely to enter the alcohol port, decreased their time spent in port, and increased their latency to enter the port.

## RESULTS

### Rats learned to discriminate between the CS+ and CS-

Viral expression and the location of optic fiber tips within the VP was confirmed for all subjects included in the analysis (Figure 1). During Pavlovian conditioning rats learned to discriminate between the alcohol-predictive CS+ and the unpaired CS-cue (Figure 2A). Port entry probability increased across days in a manner that was cue-dependent (main effect of day, F(1,402) = 15.59, p < 0.001; day by cue, F(1,402) = 75.88, p < 0.001), selectively increasing in response to the CS+ but not the CS- (Figure 2B). We did not find any significant effect of viral group on this trajectory (main effect of virus, F(1,402) = 1.43, p = 0.23; day by cue by virus, F(1,402) = 2.38, p = 0.12) and the viral groups did not differ on the final day of training prior to optogenetic manipulations (Figure 2C, main effect of virus, F(1,18) = 0.25, p = 0.62; virus by cue type, F(1,18) = 0.35, p = 0.56).

**Figure 1.**
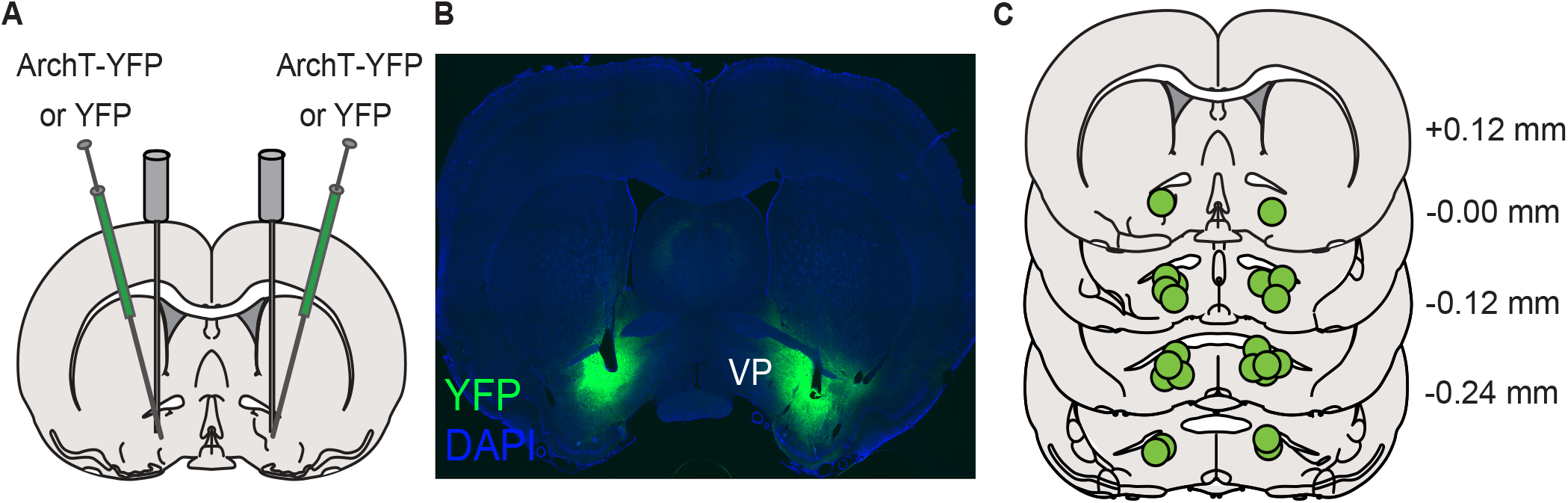
Viral approach and targeting. A) AAVs coding for the expression of ArchT-YFP or YFP alone were bilaterally infused into the ventral pallidum (VP) and optic fibers were cemented in place over the injection sites. B) Representative YFP expression localized to the VP. C) Summary of optic fiber tip locations within the VP.

**Figure 2.**
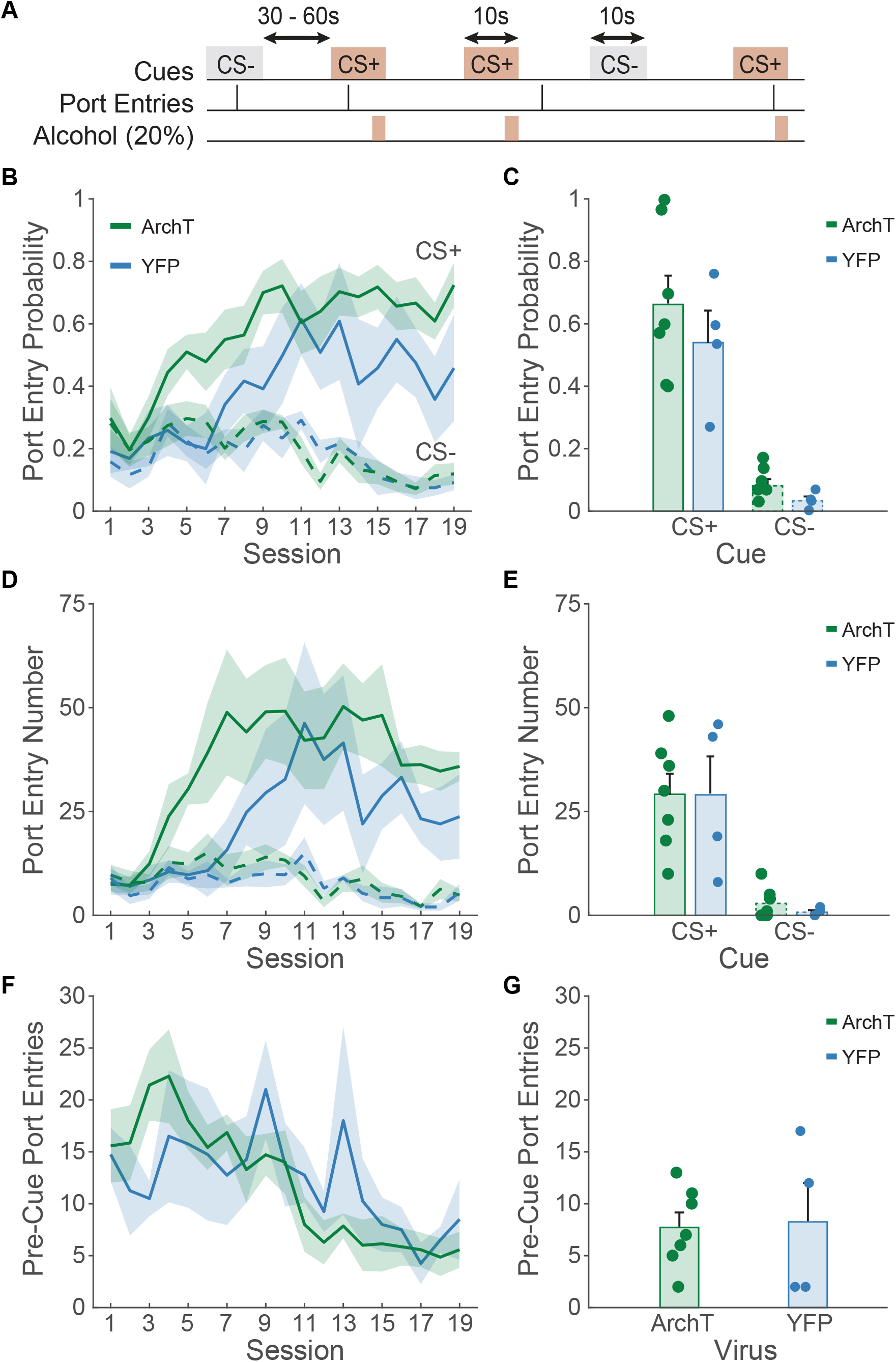
Pavlovian alcohol seeking training data. A) Schematic of the Pavlovian conditioned alcohol seeking task, where a CS+ cue predicts alcohol delivery into a reward port, and a CS-cue has no associated outcomes. B, D) Across training acquisition, rats in the ArchT and GYFP-only viral groups learned to discriminate the alcohol predictive CS+ from the CS-. C) Port entry probability and E) number of port entries during each cue type reached equal levels between the viral groups by the end of training. F) Pre-cue port entries decreased for both viral groups, and G) were equal by the end of training.

Similarly, port entry number increased in a manner that was cue-dependent (main effect of day, F(1,410) = 2.59, p = 0.10, day by cue, F(1,410) = 25.92, p < 0.001), selectively increasing during the CS+ and not the CS- (Figure 2D). We found no difference between the viral groups in their training trajectory (main effect of virus, F(1,410) = 0.24, p = 0.63; day by cue by virus, F(1,410) = 0.068, p = 0.79) and the viral groups did not differ on their final day of training prior to optogenetic manipulations (Figure 2E, main effect of virus, F(1,18) = 0.116, p = 0.74; virus by cue, F(1,18) = 0.068, p = 0.79). Port entries during the 10-sec pre-cue period decreased over time (Figure 2F, main effect of day, F(1,205) = 61.72). While we observed an interaction between day and viral group for pre-cue port entries (day by virus, F(1,205) = 6.26, p = 0.013), both groups significantly decreased their pre-cue port entry behavior (ArchT: main effect of day, F(1,131) = 71.13, p < 0.001; YFP: main effect of day, F(1,74) = 6.38, p = 0.013), and the groups did not differ in their pre-cue port entry behavior by the end of training (Figure 2G, t(9) = -0.23, p = 0.82).

By the end of training, rats also entered the port at a shorter latency during the CS+ than the CS- (main effect of cue, F(1,18) = 50.55, p < 0.001) regardless of viral group (virus by cue type, F(1,18) = 0.19, p = 0.66). They also spent more time in the port during the CS+ than the CS- (main effect of cue, F(1,18) = 12.69, p = 0.002), regardless of viral group (cue by virus, F(1,18) = 2.31, p = 0.145). Overall rats in both viral groups showed robust cue discrimination by the end of training and prior to optogenetic manipulations.

### Optogenetic inhibition of ventral pallidum disrupts Pavlovian-conditioned alcohol seeking

Following Pavlovian conditioning we assessed the effects of optogenetic inhibition of VP during 50% of CS+ and CS-presentations, pseudorandomly selected throughout the session (Figure 3A). We found that VP inhibition disrupted Pavlovian alcohol seeking according to multiple metrics.

**Figure 3.**
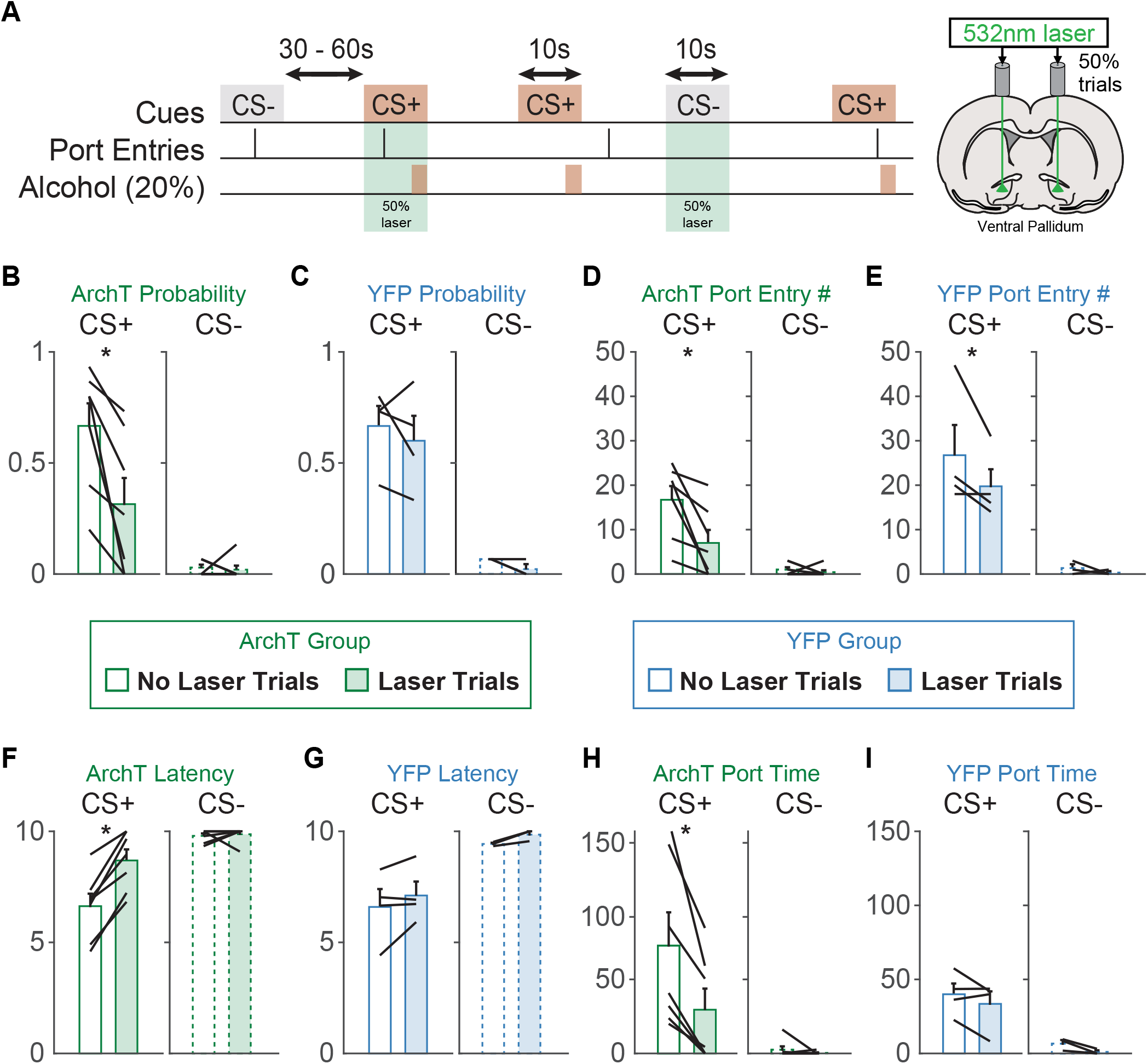
Ventral pallidum inhibition acutely disrupts Pavlovian conditioned alcohol seeking. A) Schematic of the optogenetic manipulation. Green laser was delivered on 50% of CS+ and CS-trials. B) Port entry probability during the CS+ was reduced in the ArchT group on laser stimulation trials. C) Port entry probability was not affected by laser delivery in the YFP group. D) The number of port entries during the CS+ was reduced on laser trials for the ArchT group. E) The number of port entries during the CS+ was reduced on laser trials for the YFP group. F) The latency to make port entries during the CS+ was increased on laser trials for the ArchT group. G) There was no effect of laser delivery on port entry latency for the YFP group. H) The time rats spent in the alcohol port during the CS+ was also reduced during laser trials in the ArchT group. I) There was impact of laser on port entry time for the YFP group.

First, we found that optogenetic inhibition in rats expressing ArchT in VP neurons suppressed port entry probability (main effect of laser, F(1,48) = 22.24, p < 0.001) specifically during the CS+ (Figure 3B; cue by laser, F(1,32) = p = 0.0057; main effect of cue, F(1,32) = 67.05, p < 0.001). The effect of laser depended on viral group (viral group by laser, F(1,48) = 6.04, p = 0.017; viral group by cue by laser, F(1,48) = 3.97, p = 0.051). There was no effect of laser in rats expressing YFP alone (Figure 3C, main effect of laser, F(1,16) = 0.39, p = 0.53; cue by laser, F(1,16) = <0.1, p = 0.996) who maintained robust cue discrimination (F(1,16) = 65.396, p < 0.001).

We observed less-specific effects on port entry number. Optogenetic inhibition of rats expressing ArchT in VP neurons suppressed port entry number (main effect of laser, F(1,32) = 21.52, p < 0.001) specifically during the CS+ (Figure 3D; cue by laser, F(1,32) = 10.25, p = 0.003). Unlike the effect on port entry probability, this effect did not depend on viral group (viral group by laser, F(1,48) = 1.07, p = 0.30; viral group by cue by laser, F(1,48) = 0.75, p = 0.39). We also saw significant suppression of port entry number during the CS+ in the YFP rats (Figure 3E; main effect of laser, F(1,16) = 12.83, p = 0.002; cue by laser, F(1,16) = 4.86, p = 0.04), though this appeared to be driven by one rat whose port entries on non-laser trials were >2.7 standard deviations above the mean.

Optogenetic inhibition of VP selectively altered port entry latency and time spent in port during the CS+. In rats expressing ArchT in VP optogenetic inhibition increased port entry latency (main effect of laser, F(1,32) = 29.338, p < 0.001), specifically during the CS+ and not the CS-(Figure 3F; cue by laser, F(1,32) = 13.96, p < 0.001). This effect depended on viral group (laser by viral group, F(1,48) = 7.22, p = 0.009; viral group by cue by laser, F(1,48) = 5.42, p = 0.024). Laser did not significantly impact port entry latency in rats expressing YFP-alone in VP (Figure3 G, main effect of laser, F(1,16) = 3.14, p = 0.095; cue by laser, F(1,16) = 0.11, p = 0.74). In rats expressing ArchT in VP optogenetic inhibition decreased time spent in port (main effect of laser, F(1,32) = 9.64, p = 0.0039) selectively during the CS+ and not CS-(Figure 3H; cue by laser, F(1,32) = 4.14, p = 0.050). We observed a trend towards an effect of viral group on this effect (laser by viral group, F(1,48) = 3.57, p = 0.064; viral group by cue by laser, F(1,48) = 1.85, p = 0.18) and no significant effects of laser on port time in rats expressing YFP-alone (Figure 3I, main effect of laser, F(1,16) = 2.27, p = 0.15; cue by laser, F(1,16) = 0.25, p = 0.62). Importantly, we observed no differences between the viral groups in cue-driven behavior on trials without laser. Altogether, these results suggest that optogenetic inhibition of VP selectively suppresses conditioned seeking in response to a Pavlovian alcohol predictive cue but does not disrupt the learned value of that cue or motivation for alcohol in the absence of acute VP inhibition.

## DISCUSSION

Here we find that optogenetic inhibition of VP during presentations of Pavlovian alcohol cues acutely disrupts behavioral responses to these cues. Optogenetic inhibition selectively reduces the likelihood of conditioned responses, increases the latency of these responses, and decreases time spent in the alcohol port, during the alcohol cue but not the control cue. Behavioral responses on non-laser trials do not differ between rats expressing the opsin and control rats, suggesting that VP inhibition results in an acute effect on motivated responding, and does not affect the learned or predictive value of the cues themselves on subsequent trials.

These findings are consistent with prior work showing that optogenetic inhibition of VP in general, or VP GABAergic neurons specifically, disrupts behavioral responses to a discriminative stimulus (Richard et al., 2016; Scott et al., 2023) and that optogenetic inhibition of VP disrupts Pavlovian responses to a salt-paired context (Chang et al., 2017). It remains to be seen whether overlapping populations of VP neurons are responsible for these effects, or whether more specific populations of VP neurons belonging to specific output pathways or cell-type are responsible for specific types of cue responses (Prasad et al., 2020), or seeking of rewards belonging to distinct classes (i.e. drug classes or drug versus food reward). While VP inputs to the ventral tegmental area are necessary for cue-induced reinstatement of cocaine seeking (Mahler et al., 2014), we previously found that altering the activity of VP neurons projecting to the VTA does not affect sucrose seeking in response to a discriminative stimulus (Palmer et al., 2024).

Cues predicting distinct rewards do appear to elicit distinct patterns of ensemble activity in VP (Ottenheimer et al., 2018, 2019). Previously we reported that machine learning models trained on the activity of VP neurons and ensembles in response to sucrose cues or discriminative stimuli predicting alcohol availability can reliably predict whether rats are being presented with the reward-predictive cue or the control cue. In contrast, models trained on VP responses to Pavlovian alcohol cues perform significantly worse. Importantly, this poor encoding may be partially due to the effects of non-associative alcohol exposure, which also disrupts VP encoding of Pavlovian sucrose cues. Yet, despite this poor encoding of Pavlovian alcohol cues by VP single unit and ensemble activity, we still find that normal VP activity is necessary for behavioral responses to Pavlovian alcohol cues.

### Limitations of the study

While our results suggest the normal VP activity is critical for Pavlovian conditioned alcohol-seeking, our results do not address whether cue-evoked changes in activity are responsible for these behaviors, or whether basal VP activity is necessary. Our optogenetic manipulation presumably inhibited activity below basal levels, in addition to disrupting cue-evoked excitations. Additionally, while we have observed cue-evoked citations in approximately 30% of neurons, almost 20% of neurons are inhibited by Pavlovian alcohol cues (Ottenheimer et al., 2019). These excited and inhibited neurons may constitute subsets of GABAergic and glutamatergic VP neurons which have been shown to have opposing responses to appetitive and aversive stimuli, and to drive oppositely-valenced behaviors (Faget et al., 2018; Tooley et al., 2018; Heinsbroek et al., 2020; Stephenson-Jones et al., 2020). Here we did not target our optogenetic manipulations based on cell-type, though the correspondence between the aforementioned activity patterns during Pavlovian alcohol-seeking and previously defined cell-types is not yet established.

Here we only assessed the impact of optogenetic inhibition on Pavlovian alcohol-seeking in male rats. A rich prior literature, including our own work, has documented that female rodents, including Long Evans rats, consume more g/kg ethanol on average than male rodents, under both continuous and intermittent access conditions (Li and Lumeng, 1984; Lancaster and Spiegel, 1992; Juárez and De Tomasi, 1999; Priddy et al., 2017; Lorrai et al., 2019; Aguirre et al., 2020; Carpio et al., 2022; Armstrong et al., 2023). While we previously reported no significant sex differences in training trajectories or plateaus during both Pavlovian conditioning for alcohol and training with an alcohol discriminative stimulus (Carpio et al., 2022; Armstrong et al., 2023), it is possible that Pavlovian conditioned responses to alcohol cues in female rats may be differentially impacted by, or resistant to, peri-cue inhibition of the VP.

### Conclusions and future directions

Altogether we find that disruptions to VP activity during alcohol cue presentations disrupt behavioral responses to these cues. Future work should interrogate whether VP activity is necessary for behavioral responses to alcohol cues in models of relapse. Additionally, limited work has been done to establish neural mechanisms by which cues for alcohol and other drugs can elicit compulsive drug and alcohol seeking that is resistant to negative consequences. Importantly, VP activity has been implicated in reward-seeking under conditions of conflict or risk (Farrell et al., 2021). Distinct populations of VP neurons, as well as input and output pathways, likely underly the ability of cues to promote craving, and disrupt inhibitory control or sensitivity to negative outcomes or punishment. Experiments targeting more specific VP populations during models of cue-elicited alcohol seeking may reveal more nuanced functions of VP in cue-elicited alcohol-seeking.

## METHODS

### Subjects

Male Long Evans rats (n=11; Envigo) weighing 250-275 grams at arrival served as experimental subjects and were individually housed in a temperature- and humidity-controlled colony room on a 12 h light/dark cycle. Rats were fed ad libitum, and water was provided ad libitum to all rats. All experimental procedures were approved by the Institutional Animal Care and Use Committees at Johns Hopkins University and the University of Minnesota and were carried out in accordance with the guidelines on animal care and use of the National Institutes of Health of the United States.

### Alcohol pre-exposure

Prior to surgery and Pavlovian conditioning rats were pre-exposed to ethanol in the home cage as described previously (Simms et al., 2008; Remedios et al., 2014; Millan et al., 2017; Armstrong et al., 2023). First, they received 1 week of continuous access to 10% ethanol, followed by intermittent access to 20% ethanol for 24 hours at a time, starting on Monday, Wednesday and Friday (MWF), for a period of 7 weeks. Rats that consumed more than 2 g/kg per day during the last 2 weeks of pre-exposure underwent surgery for viral infusions and fiber implantation prior to Pavlovian conditioning.

### Surgeries

Following alcohol pre-exposure rats received infusions of virus and optical fiber implants for optogenetic inhibition of VP. During surgery, rats were anesthetized with isoflurane (5%) and placed in a stereotaxic apparatus, after which surgical anesthesia was maintained with isoflurane (0.5–2.0%). Rats received preoperative injections of carprofen (5 mg/kg) for analgesia and cefazolin (75 mg/kg) to prevent infection. First, 0.5 μl of virus containing the archaerhodopsin gene construct (n=7; AAV5/ CamKIIa-ArchT3.0-eYFP, 7x1012 viral particles/ml from the University of North Carolina Vector Core) or control virus (n=4; AAV5-CamKIIa-eYFP) was delivered bilaterally to VP through 28-gauge injectors at a rate of 0.1 μl per min for 5 minutes. Injectors were left in place for 10 min following the infusion to allow virus to diffuse away from the infusion site. Injector tips were aimed at the following coordinates in relation to Bregma: 0.0 mm AP, +/-2.5 mm mediolateral, -8.2 mm dorsoventral. Then, rats were implanted with 300-micron diameter optic fibers aimed .3 mm above the center of the virus infusion. Implants were secured to the skull with bone screws and dental acrylic. Rats recovered for at least one week before any behavioral training. To allow for sufficient viral expression, rats recovered for at least 4 weeks before optogenetic testing manipulations.

### Pavlovian conditioning with alcohol

Prior to cue conditioning, rats underwent port training in which they received 30 deliveries of .065 mL 20% ethanol to a reward port. Next, they underwent Pavlovian conditioning for at least 19 sessions (MWF) prior to optogenetic manipulations as described previously (Ottenheimer et al., 2019). Rats were randomly assigned one of the following cues as their conditioned stimulus (CS+) for training and testing: white noise or a 2900 Hz tone. Rats received the alternate auditory cue as their CS-. During each conditioning sessions, the CS+ and CS-, each lasting 10s, were presented 30 times on a pseudo-random variable interval schedule with a mean inter-trial interval (ITI) of 80s. At 9s after the CS+ onset, 0.065 mL of 20% ethanol was delivered into the reward delivery port over a period of 1s. Rats were initially trained untethered to optical testing cables and were tethered starting on session 11.

Optogenetic inhibition during Pavlovian alcohol-seeking Rats were tested while tethered to optical patch cables (200 μm core, 0.4 m split cables, Doric), connected to a rotary join on a cantilever arm, which was connected to a 532 nm DPSS laser. On test days, rats received continuous bilateral photoillumination of the VP (10-15 mW light power) on 50% of CS+ and CS-trials, pseudo-randomly selected. Photoillumination was initiated at the start of the cue and terminated at the end of the cue.

### Histological assessment

Following the completion of behavioral testing, rats were anesthetized with pentobarbital and perfused intracardially with 9% saline followed by 4% paraformaldehyde. Brains were extracted, post-fixed in 4% paraformaldehyde for up to 24 hours, and placed in 25% sucrose for at least 48 hours, before being sectioned at 50 μm. Sections were then mounted, coverslipped with VEC-TASHIELD mounting medium with DAPI, and imaged on a Zeiss Axio 2 microscope to confirm the presence of viral expression in VP. A subset of tissue was immunostained for GFP using mouse anti-GFP (1:1500, Invitrogen #A11120) and Alexa Fluor 488 donkey anti mouse (1:500, Invitrogen #A21202). Sections were washed in PBS with Triton (PBST) 3 times for 5 minutes, incubated in 10% normal donkey serum in PBST for 60 minutes, and then incubated in primary antibody solution in PBST overnight at 4C. Sections were then washed in PBST 3 times and incubated in secondary antibody solution in PBST for 2 hours. Following three additional washes in PBST, sections were mounted, coverslipped with VEC-TASHIELD mounting medium with DAPI and imaged on a Nikon Ti-E Inverted Deconvolution Microscope (University Imaging Centers, University of Minnesota). Location of optic fiber tips in the ventral pallidum was confirmed and mapped onto a standardized rat brain atlas (Figure 1).

### Statistical Analysis

Our primary behavioral measure during Pavlovian conditioning and testing sessions was the number of port entries during each epoch (CS+, CS- and/or ITI) and port entry probability during each cue (i.e. the proportion of trials with a port entry). We also assessed port entry latency and time spent in port on optogenetic manipulation days. To account for a mixture of within-subject and between-subjects data collection we analyzed the data using linear mixed effects models (fitlme function with maximum likelihood estimation) in MATLAB (Mathworks). To assess the development of cue discrimination across Pavlovian conditioning sessions, we fit models with fixed effects for cue type and session and a random effect for subject. To assess the impact of optogenetic inhibition on responses to the cues we fit models with fixed effects for cue type, laser status, and virus and a random effect for subject.

## Funding and Acknowledgements

This work was supported in part by National Institutes of Health grants K99AA025384 and R00AA025384 to JMR and R01AA027213 to PHJ.

## REFERENCES

Aguirre CG, Stolyarova A, Das K, Kolli S, Marty V, Ray L, Spigelman I, Izquierdo A (2020) Sex-dependent effects of chronic intermittent voluntary alcohol consumption on attentional, not motivational, measures during probabilistic learning and reversal. PLOS ONE 15:e0234729.

Armstrong A, Rosenthal H, Stout N, Richard JM (2023) Reinstatement of Pavlovian responses to alcohol cues by stress. Psychopharmacology 240:531–545.

Carpio MJ, Gao R, Wooner E, Cayton CA, Richard JM (2022) Alcohol availability during withdrawal gates the impact of alcohol vapor exposure on responses to alcohol cues. Psychopharmacology Available at: https://pubmed.ncbi.nlm.nih.gov/35881146/ [Accessed September 11, 2022].

Chang SE, Smedley EB, Stansfield KJ, Stott JJ, Smith KS (2017) Optogenetic inhibition of ventral pallidum neurons impairs context-driven salt-seeking. Journal of Neuroscience 37:5670–5680.

de Wit H (2009) Impulsivity as a determinant and consequence of drug use: a review of underlying processes. Addict Biol 14:22–31.

Everitt BJ, Robbins TW (2005) Neural systems of reinforcement for drug addiction: from actions to habits to compulsion. Nat Neurosci 8:1481–1489.

Faget L, Zell V, Souter E, McPherson A, Ressler R, Gutierrez-Reed N, Yoo JH, Dulcis D, Hnasko TS (2018) Opponent control of behavioral reinforcement by inhibitory and excitatory projections from the ventral pallidum. Nature Communications 9:849.

Farrell MR, Esteban JSD, Faget L, Floresco SB, Hnasko TS, Mahler SV (2021) Ventral pallidum GABA neurons mediate motivation underlying risky choice. Journal of Neuroscience 41:4500–4513.

Farrell MR, Ruiz CM, Castillo E, Faget L, Khanbijian C, Liu S, Schoch H, Rojas G, Huerta MY, Hnasko TS, Mahler SV (2019) Ventral pallidum is essential for cocaine relapse after voluntary abstinence in rats. Neuropsychopharmacology: 1–12.

Farrell MR, Ye Q, Xie Y, Esteban JSD, Mahler SV (2022) Ventral pallidum GABA neurons bidirectionally control opioid relapse across rat behavioral models. Addict Neurosci 3:100026.

Field M, Jones A (2017) Elevated alcohol consumption following alcohol cue exposure is partially mediated by reduced inhibitory control and increased craving. Psychopharmacology 234:2979–2988.

Garbusow M, Sebold M, Beck A, Heinz A (2014) Too Difficult to Stop: Mechanisms Facilitating Relapse in Alcohol Dependence. Neuropsychobiology 70:103–110.

Garland EL, Franken IHA, Howard MO (2012) Cue-elicited heart rate variability and attentional bias predict alcohol relapse following treatment. Psychopharmacology 222:17–26.

Heinsbroek JA, Bobadilla AC, Dereschewitz E, Assali A, Chalhoub RM, Cowan CW, Kalivas PW (2020) Opposing Regulation of Cocaine Seeking by Glutamate and GABA Neurons in the Ventral Pallidum. Cell Reports 30:2018-2027.e3.

Jones A, Rose AK, Cole J, Field M (2013) Effects of Alcohol Cues on Craving and Ad Libitum Alcohol Consumption in Social Drinkers: The Role of Disinhibition. Journal of Experimental Psychopathology 4:239–249.

Juárez J, De Tomasi EB (1999) Sex Differences in Alcohol Drinking Patterns During Forced and Voluntary Consumption in Rats. Alcohol 19:15–22.

Kalivas PW, Volkow ND (2005) The neural basis of addiction: a pathology of motivation and choice. Am J Psychiatry 162:1403–1413.

Koordeman R, Anschutz DJ, Engels RCME (2011) Exposure to alcohol commercials in movie theaters affects actual alcohol consumption in young adult high weekly drinkers: an experimental study. Am J Addict 20:285–291.

Kupchik YM, Prasad AA (2021) Ventral pallidum cellular and pathway specificity in drug seeking. Neurosci Biobehav Rev 131:373–386.

Lancaster FE, Spiegel KS (1992) Sex differences in pattern of drinking. Alcohol 9:415–420.

Lederman J, Lardeux S, Nicola SM (2021) Vigor Encoding in the Ventral Pallidum. eneuro 8:ENEURO.0064-21.2021.

Li T-K, Lumeng L (1984) Alcohol Preference and Voluntary Alcohol Intakes of Inbred Rat Strains and the National Institutes of Health Heterogeneous Stock of Rats. Alcoholism: Clinical and Experimental Research 8:485–486.

Lorrai I, Contini A, Gessa GL, Mugnaini C, Corelli F, Colombo G, Maccioni P (2019) Operant, oral alcohol self-administration: Sex differences in Sardinian alcohol-preferring rats. Alcohol 79:147–162.

Mahler SV, Vazey EM, Beckley JT, Keistler CR, McGlinchey EM, Kaufling J, Wilson SP, Deisseroth K, Woodward JJ, Aston-Jones G (2014) Designer receptors show role for ventral pallidum input to ventral tegmental area in cocaine seeking. Nature neuroscience 17:577–585.

McFarland K, Davidge SB, Lapish CC, Kalivas PW (2004) Limbic and motor circuitry underlying footshock-induced reinstatement of cocaine-seeking behavior. The Journal of neuroscience : the official journal of the Society for Neuroscience 24:1551–1560.

McFarland K, Kalivas PW (2001) The circuitry mediating cocaine-induced reinstatement of drug-seeking behavior. The Journal of neuroscience : the official journal of the Society for Neuroscience 21:8655–8663.

Millan EZ, Kim HA, Janak PH (2017) Optogenetic activation of amygdala projections to nucleus accumbens can arrest conditioned and unconditioned alcohol consummatory behavior. Neuroscience Available at: http://www.ncbi.nlm.nih.gov/pubmed/28757250 [Accessed August 3, 2017].

Niaura RS, Rohsenow DJ, Binkoff JA, Monti PM, Pedraza M, Abrams DB (1988) Relevance of cue reactivity to understanding alcohol and smoking relapse. Journal of Abnormal Psychology 97:133–152.

Ottenheimer D, Richard JM, Janak PH (2018) Ventral pallidum encodes relative reward value earlier and more robustly than nucleus accumbens. Nature Communications 9:4350.

Ottenheimer DJ, Wang K, Haimbaugh A, Janak PH, Richard JM (2019) Recruitment and disruption of ventral pallidal cue encoding during alcohol seeking. European Journal of Neuroscience 50:3428–3444.

Palmer D, Cayton CA, Scott A, Lin I, Newell B, Paulson A, Weberg M, Richard JM (2024) Ventral pallidum neurons projecting to the ventral tegmental area reinforce but do not invigorate reward-seeking behavior. Cell Reports 43:113669.

Perry CJ, McNally GP (2013) A role for the ventral pallidum in context-induced and primed reinstatement of alcohol seeking. The European journal of neuroscience 38:2762–2773.

Poisson CL, Engel L, Saunders BT (2021) Dopamine Circuit Mechanisms of Addiction-Like Behaviors. Front Neural Circuits 15:752420.

Prasad AA, McNally GP (2016) Ventral Pallidum Output Pathways in Context-Induced Reinstatement of Alcohol Seeking. Journal of Neuroscience 36:11716–11726.

Prasad AA, Xie C, Chaichim C, Nguyen JH, McClusky HE, Killcross S, Power JM, McNally GP (2020) Complementary roles for ventral pallidum cell types and their projections in relapse. The Journal of Neuroscience:0262–19.

Priddy BM, Carmack SA, Thomas LC, Vendruscolo JCM, Koob GF, Vendruscolo LF (2017) Sex, strain, and estrous cycle influences on alcohol drinking in rats. Pharmacology Biochemistry and Behavior 152:61–67.

Remedios J, Woods C, Tardif C, Janak PH, Chaudhri N (2014) Pavlovian-conditioned alcohol-seeking behavior in rats is invigorated by the interaction between discrete and contextual alcohol cues: implications for relapse. Brain and behavior 4:278–289.

Richard JM, Ambroggi F, Janak PH, Fields HL (2016) Ventral Pallidum Neurons Encode Incentive Value and Promote Cue-Elicited Instrumental Actions. Neuron 90:1165–1173.

Richard JM, Fields HL (2016) Mu-opioid receptor activation in the medial shell of nucleus accumbens promotes alcohol consumption, self-administration and cue-induced reinstatement. Neuropharmacology 108:14–23.

Richard JM, Stout N, Acs D, Janak PH (2018) Ventral pallidal encoding of reward-seeking behavior depends on the underlying associative structure. eLife 7:e33107.

Robinson TE, Berridge KC (2004) Incentive-sensitization and drug “wanting.” Psychopharmacology (Berl) 171:352–353.

Root DH, Melendez RI, Zaborszky L, Napier TC (2015) The ventral pallidum: Subregion-specific functional anatomy and roles in motivated behaviors. Progress in neurobiology 130:29–70.

Saunders BT, Richard JM, Janak PH (2015) Contemporary approaches to neural circuit manipulation and mapping: focus on reward and addiction. Philosophical transactions of the Royal Society of London Series B, Biological sciences 370 Available at: http://www.ncbi.nlm.nih.gov/pubmed/26240425 [Accessed August 5, 2015].

Sayette MA, Hufford MR (1994) Effects of cue exposure and deprivation on cognitive resources in smokers. J Abnorm Psychol 103:812–818.

Scott A, Palmer D, Newell B, Lin I, Cayton CA, Paulson A, Remde P, Richard JM (2023) Ventral Pallidal GABAergic Neuron Calcium Activity Encodes Cue-Driven Reward Seeking and Persists in the Absence of Reward Delivery. Journal of Neuroscience 43:5191–5203.

Shaham Y, Shalev U, Lu L, de Wit H, Stewart J (2003) The reinstatement model of drug relapse: history, methodology and major findings. Psychopharmacology 168:3–20.

Simms JA, Steensland P, Medina B, Abernathy KE, Chandler LJ, Wise R, Bartlett SE (2008) Intermittent access to 20% ethanol induces high ethanol consumption in Long-Evans and Wistar rats. Alcoholism, clinical and experimental research 32:1816–1823.

Smith KS, Tindell AJ, Aldridge JW, Berridge KC (2009) Ventral pallidum roles in reward and motivation. Behavioural brain research 196:155–167.

Soares-Cunha C, Heinsbroek JA (2023) Ventral pallidal regulation of motivated behaviors and reinforcement. Front Neural Circuits 17:1086053.

Stein KD, Goldman MS, Del Boca FK (2000) The influence of alcohol expectancy priming and mood manipulation on subsequent alcohol consumption. J Abnorm Psychol 109:106–115.

Stephenson-Jones M, Bravo-Rivera C, Ahrens S, Furlan A, Xiao X, Fernandes-Henriques C, Li B (2020) Opposing Contributions of GABAergic and Glutamatergic Ventral Pallidal Neurons to Motivational Behaviors. Neuron Available at: https://linkinghub.elsevier.com/retrieve/pii/S0896627319310487 [Accessed January 20, 2020].

Stewart J, de Wit H, Eikelboom R (1984) Role of unconditioned and conditioned drug effects in the self- administration of opiates and stimulants. Psychological Review 91:251–268.

Tang X-C, McFarland K, Cagle S, Kalivas PW (2005) Cocaine-Induced Reinstatement Requires Endogenous Stimulation of {micro}-Opioid Receptors in the Ventral Pallidum. J Neurosci 25:4512–4520.

Tiffany ST, Carter BL (1998) Is craving the source of compulsive drug use? Journal of Psychopharmacology 12:23–30.

Tindell AJ, Berridge KC, Aldridge JW (2004) Ventral pallidal representation of pavlovian cues and reward: population and rate codes. J Neurosci 24:1058–1069.

Tindell AJ, Berridge KC, Zhang J, Peciña S, Aldridge JW (2005) Ventral pallidal neurons code incentive motivation: amplification by mesolimbic sensitization and amphetamine. Eur J Neurosci 22:2617–2634.

Tooley J, Marconi L, Alipio JB, Matikainen-Ankney B, Georgiou P, Kravitz AV, Creed MC (2018) Glutamatergic Ventral Pallidal Neurons Modulate Activity of the Habenula–Tegmental Circuitry and Constrain Reward Seeking. Biological Psychiatry 83:1012–1023.

Tsiang MT, Janak PH (2006) Alcohol seeking in C57BL/6 mice induced by conditioned cues and contexts in the extinction-reinstatement model. Alcohol (Fayetteville, NY) 38:81–88.

Valyear MD, Villaruel FR, Chaudhri N (2017) Alcohol-seeking and relapse: A focus on incentive salience and contextual conditioning. Behavioural Processes 141:26–32.

